# The Fontan EV Score: A Circulating Extracellular Vesicle-Based Risk Stratification Tool for Fontan-Associated Liver Disease

**DOI:** 10.64898/2026.07.31.742169

**Authors:** Felipe Takaesu, Xinlei Li, Jennifer Kievert, Ashley Zhou, Sherri Kemper, Satoshi Yuhara, Syed Faizullah Hussaini, Tatsuya Watanabe, Junya Matsuda, Fahd Taha, Adrienne Morrison, Kirsten Nelson, Jocelyhnn Zucco, Aymer Naguib, Christopher McKee, Jordan Hill, Sergio A. Carrillo, Christopher K. Breuer, John M. Kelly, David R. Brigstock, Michael E. Davis

## Abstract

**Background:** Fontan-associated liver disease (FALD) is a universal complication of the Fontan palliation characterized by chronic congestion and progressive hepatic fibrosis. Current diagnostics rely on invasive biopsies or non-specific biochemical and imaging biomarkers that fail to capture early fibrogenesis, creating a critical need for non-invasive biomarkers to stratify disease severity.

**Methods:** We utilized a translational ovine Fontan model (n = 19) to investigate circulating serum extracellular vesicles (sEVs) as reporters of hepatic pathology. Longitudinal serum samples paired with liver elastography were collected, and sEVs were subjected to multi-omic profiling including small RNA sequencing and proteomics. Regularized regression was used to identify transcriptomic predictors, which were integrated with time post-surgery into an ordinal logistic regression framework to construct the Fontan EV Score (FES). Model performance was evaluated on a held-out test cohort and benchmarked against established serological fibrosis indices. To validate the biological relevance of the FES panel, TGF-β-treated human liver organoids were generated and scored miRNA expression was assessed.

**Results:** The sEV proteome exhibited robust separation by surgical physiology, while the small RNA cargo was primarily stratified by fibrotic status. Bioinformatic analysis confirmed a high hepatic origin for these transcripts and identified enrichment of inflammatory pathways including Toll-like receptor and Interleukin-17 cascades in fibrotic subjects. The FES, incorporating time post-surgery and eleven small RNA biomarkers, demonstrated high predictive accuracy in the independent testing cohort with an AUC of 0.876 for moderate and 0.963 for severe fibrosis, substantially outperforming APRI (AUC = 0.618) and FIB-4 (AUC = 0.731). In TGF-β-treated human liver organoids, several scoring miRNAs, including miR-125a-5p and miR-193b-5p, were directionally responsive to profibrotic stimulation.

**Conclusions:** Circulating sEVs carry a liver-associated molecular cargo that can be leveraged for the non-invasive prediction of FALD severity. The FES provides a biologically validated scoring system that substantially outperforms existing serological indices and offers a new avenue for early detection and risk stratification of FALD

**Novelty and Significance:** *What is Known?:*

- Fontan-associated liver disease (FALD) is a nearly universal consequence of the Fontan circulation, driven by chronic venous hypertension and reduced cardiac output.
- Current surveillance tools, including transaminases, composite serological indices (APRI, FIB-4), and elastography, have limited sensitivity and specificity for detecting and staging hepatic fibrosis in the Fontan population.
- Circulating small extracellular vesicles (sEVs) carry tissue-derived molecular cargo and have shown diagnostic potential in other liver diseases, but their utility in FALD has not been explored.

*What New Information Does This Article Contribute?:*

- Multi-omic profiling of circulating sEVs in a translational ovine Fontan model reveals that the small RNA cargo is stratified by fibrotic status and enriched for inflammatory pathways associated with hepatic stellate cell activation.
- The Fontan EV Score (FES), integrating time post-surgery with eleven circulating small RNA biomarkers, predicts FALD severity with substantially greater accuracy than APRI and FIB-4.
- TGF-β-treated human liver organoids confirm that several FES-associated miRNAs are directly responsive to profibrotic stimulation, providing biological validation independent of Fontan hemodynamics. This study demonstrates that circulating sEVs function as non-invasive reporters of hepatic fibrogenesis in the Fontan circulation and introduces the first EV-based scoring system for FALD risk stratification. The FES achieved an AUC of 0.876 for moderate and 0.963 for severe fibrosis in an independent test cohort, outperforming established serological indices that were originally developed for viral hepatitis but which perform poorly in congestive hepatopathy. By combining molecular biomarker discovery with in vitro functional validation, this work establishes a foundation for developing targeted, non-invasive diagnostics to guide surveillance and clinical decision-making in the growing Fontan patient population.

## Introduction

Fontan palliation is the definitive treatment for patients born with single ventricle heart disease. While lifesaving, it is a palliative treatment, with significant late term morbidity and reduced duration of life. Among the most pressing long-term complications are Fontan-associated liver disease (FALD)^1,2^. As survival rates following the Fontan procedure improve, the prevalence of FALD continues to rise and represents an emerging medical challenge^3^. Current literature suggests that the chronic venous hypertension and reduced cardiac output inherent to the Fontan circulation are the primary drivers of this pathology^2,3^. However, the precise mechanisms underlying the development of FALD remain unknown. This gap in knowledge is largely attributed to the difficulty in studying the clinical progression of the disease longitudinally. Consequently, determining the specific molecular causes for the progression of FALD is essential to improve our ability to diagnose and treat this serious condition.

Current clinical surveillance for FALD relies heavily on a combination of serological markers and imaging techniques; However, each possesses significant limitations^4^. Liver biopsy remains the gold standard for staging fibrosis but its invasive nature carries risks of bleeding and infection^5,6^. Furthermore, biopsy samples are limited to only evaluating a small portion of the liver, which is problematic given the heterogenous nature of the disease, and is impractical for frequent longitudinal monitoring to capture the dynamic changes that occur over time. Non-invasive alternatives such as transient elastography and magnetic resonance elastography are increasingly utilized to estimate liver stiffness^4^, but these imaging techniques can be confounded by hepatic congestion which is a hallmark of Fontan physiology^7^. Currently, the most standard biochemical liver function tests include measuring the quantity of aspartate aminotransferase (AST) and alanine aminotransferase (ALT) in serum, but recent reports highlight that both enzymes can remain within normal limits even in the presence of advanced hepatic fibrosis or cirrhosis in Fontan patients^3^. Composite serological indices such as FIB-4 and APRI, which were originally developed for viral hepatitis, have similarly shown limited accuracy in the Fontan population due to the non-inflammatory nature of congestive hepatopathy^8–10^. This disconnect renders current routine blood work insufficient for early detection and highlights the urgent need for sensitive and specific biomarkers that can accurately stratify risk before irreversible damage occurs.

Emerging studies have highlighted circulating small extracellular vesicles (sEVs) as critical mediators of intercellular communication and promising candidates for non-invasive diagnostics^11–13^. These lipid-bound nanovesicles encapsulate a rich cargo of proteins, lipids, and nucleic acids that mirrors the cytosolic composition and physiological state of their parent cell^14,15^. Consequently, circulating sEVs can be exploited as a type of liquid biopsy that allows for the investigation of tissue-specific pathology through non-invasive blood sampling. This diagnostic potential has been shown in oncology where sEV-associated microRNAs and surface proteins serve as biomarkers for tumor burden and metastatic progression^16,17^. Similarly, in the context of hepatic disease, sEVs facilitate the crosstalk between hepatocytes and non-parenchymal cells during the pathogenesis of alcoholic liver disease^18^. Because these vesicles can be isolated from serum, they present a unique opportunity to monitor the molecular progression of FALD non-invasively.

Given these previous studies, we hypothesized that FALD progression can be similarly monitored through sEV signaling, and here, we leverage a clinically relevant ovine Fontan survival model to investigate the molecular signals associated with the development of FALD. We utilized serial serum sampling to isolate sEVs and performed a comprehensive multi-omic characterization of their protein and RNA cargo over a longitudinal time course. To translate these molecular findings into a clinical tool, we employed regularized regression techniques to identify robust transcriptomic predictors which we subsequently integrated with time post-surgery using an ordinal logistic regression framework. By correlating these molecular features with longitudinal hepatic elastography data, we established a novel scoring system, the Fontan EV Score (FES), capable of stratifying the severity of FALD. We validated the FES against an independent test cohort and benchmarked its performance against established serological fibrosis indices. This work demonstrates that circulating sEV signatures can serve as robust non-invasive biomarkers to predict liver stiffness and provides a new avenue for the early detection and management of FALD.

## Methods

### Data Availability

The data that support the findings of this study are available from the corresponding author upon reasonable request. Small RNA sequencing data have been deposited in the Gene Expression Omnibus under accession number GSE[XXXXXX]. Please see the Major Resources Table in the Supplemental Materials for all materials used in this study.

### Ethical Statement

All animal studies were performed according to ARRIVE Guidelines and under the approval and guidance of the Nationwide Children’s Hospital Animal Welfare and Resource Committee (Approval AR13-00079 and AR24-00179). Sheep were housed in facilities that are USDA licensed and AAALAC accredited. Housing space for sheep was in accordance with the 2011 Guide for the Care and Use of Laboratory Animals (including HVAC parameters, lighting, and airflow).

### Establishing the Fontan Connection

Details on the development and characterization of the ovine Fontan model are available in our group’s previous work^19–22^. The animal model of Fontan hemodynamics was established in sheep with normal biventricular anatomy by detaching the inferior vena cava from the right atrium and reconnecting it to the main pulmonary artery via an end-to-end anastomosis facilitated by a polytetrafluorethylene conduit (similar to a extracardiac conduit as is performed in human patients) and the superior vena directly anastomosed to the main pulmomary artery as described previously. Model hemodynamics of systemic venous hypertension are like those seen in human Fontan patients. Following the operation, sheep were closely monitored for approximately 2 weeks in an ICU.

Abdominal ultrasound exams were performed at regular intervals using a Mindray Resona 7 ultrasound system and a SC6-1E curved array transducer. Animals were sedated with propofol, intubated for mechanical ventilation, and placed in the left lateral decubitus position. During ultrasound exams, the width of the portal and hepatic veins was measured. Color Doppler was used to record the pulse wave velocities of the portal vein, hepatic artery, and hepatic vein. The resistive index of the hepatic artery was calculated from acquired systolic and diastolic velocities. Ultrasound elastography was performed using the STQ point shear wave elastography (pSWE) function in accordance with manufacturer guidelines and clinical guidelines^23,24^. Elastography measurements were obtained in four regions; peripheral parenchyma adjacent to the inferior vena cava, central parenchyma adjacent to the portal vein, medial parenchyma and central parenchyma adjacent to the hepatic vein. For each region, 12 measurements were captured when the motion stability index (M-STB) displayed a stable state with minimal motion interference while undergoing breath-hold. Average shear wave velocity (Cs) of each region was reported when the interquartile range (IQR) to median ratio was ≤ 15%.

### Percutaneous Liver Biopsy

Ultrasound-guided percutaneous liver biopsies were performed during the same anesthetic event as the abdominal ultrasound examinations. Animals were placed in the left lateral decubitus position, and the abdominal area was sterilely prepped using alcohol and chlorhexidine. The portal vein was identified using an SC-1U transducer on the Mindray Resona 7 ultrasound system. Point shear wave elastography was performed as previously described in the region targeted for biopsy. A 16-gauge biopsy needle was inserted approximately 2–3 cm into the hepatic parenchyma adjacent to the portal vein under ultrasound guidance. Three to four core biopsies were collected per study. One biopsy core was fixed in 10% neutral-buffered formalin for two weeks, after which the formalin was replaced with 70% ethanol. Fixed samples were subsequently embedded in paraffin, sectioned, and mounted on slides by the Nationwide Children’s Hospital Histopathology Core.

### Necropsy Liver Biopsy

At the time of euthanasia, the liver was explanted and divided into right and left lobes. The right lobe was sectioned into 1-inch segments. From each segment, cross-sectional samples were obtained using a 6 mm biopsy punch, yielding cores of approximately 6 mm × 4 mm. Biopsy cores were fixed in 10% neutral-buffered formalin for two weeks, after which the formalin was replaced with 70% ethanol. Fixed samples were embedded in paraffin, sectioned, and mounted on slides by the Nationwide Children’s Hospital Histopathology Core.

### Histological Staining and Imaging

For PSR staining, formalin-fixed paraffin-embedded (FFPE) sections were deparaffinized in xylene and rehydrated through a graded ethanol series (100%, 90%, 70%) to distilled water. Sections were stained with Weigert’s iron hematoxylin (Electron Microscopy Sciences, Cat# 26044-06) for 30 minutes, followed by a 10-minute rinse in running tap water. Sections were then incubated in Picrosirius Red solution (0.1% Direct Red 80 in saturated aqueous picric acid) for 1 hour, differentiated in 0.5% acidified water (0.5% glacial acetic acid), dehydrated through graded ethanol, cleared in xylene, and cover slipped with permanent mounting medium.

For α-SMA immunohistochemistry, FFPE sections were deparaffinized and rehydrated as above. Heat-induced antigen retrieval was performed in 10 mM sodium citrate buffer (pH 6.0, 0.05% Tween 20) using a pressure cooker for 10 minutes, followed by cooling to room temperature. Endogenous peroxidase activity was quenched with 3% hydrogen peroxide for 10 minutes. Sections were blocked with Background Sniper (Biocare Medical, Cat# BS966M) supplemented with 3% normal goat serum and avidin (Avidin/Biotin Blocking Kit, Vector Laboratories, Cat# SP-2001) for 30 minutes. Sections were then incubated overnight at 4°C with monoclonal mouse anti-human smooth muscle actin (clone 1A4, Agilent, Cat # M085101-2) diluted 1:2000 in 0.1% BSA-PBS with biotin blocking solution. Following washes in PBS-Tween, sections were incubated with biotinylated goat anti-mouse IgG secondary antibody (Vector Laboratories, Cat# BA-9200-1.5) at 1:1500 dilution in 0.1% BSA-PBS for 30 minutes at room temperature. Signal amplification was performed using VECTASTAIN Elite ABC Reagent (Vector Laboratories, Cat# PK-7100) for 30 minutes, and chromogenic detection was carried out with ImmPACT DAB Substrate Kit (Vector Laboratories, Cat# SK-4105) for 2 minutes. Sections were counterstained with Gill’s hematoxylin for 2 minutes, blued in Scott’s solution (0.2% sodium bicarbonate, 1% magnesium sulfate) for 10 minutes, dehydrated, cleared, and coverslipped with VectaMount permanent mounting medium.

All stained slides were imaged using a Zeiss Observer.Z1 inverted microscope at 10× magnification with tile scanning to capture the full biopsy cross-section.

### Blood Sample Collection and Extracellular Vesicle Characterization

For both the Fontan cohort and surgical control cohort, blood samples were collected daily from the date of surgery to the end of each animal’s patient-specific ICU monitoring endpoint. Collected blood samples were utilized for ICU monitoring and for serum proteomic analysis. Samples were harvested into a serum or plasma blood collection tube (BD Vacutainer, Cat # 02-683-94 and BD Vacutainer, Cat # 362788, respectively) for 15 min at room temperature. Samples were then spun at 1500 × g for 15 min at 4 °C followed by immediate serum or plasma collection from the supernatant. Samples were stored at -80 °C for long-term use.

Serum and plasma samples were thawed on ice in preparation for sEV isolation. 1000 µL of sample were centrifuged at 10,000xg for 10 minutes to remove large particles and debris. 500 µL of supernatant were loaded into a qEV 35nm original column (Izon Science, Christchurch, New Zealand, Cat # ICO-35) to isolate sEVs. sEV fractions were eluted in 1 mL of PBS and stored at -80°C for downstream applications. sEV size and concentration were measured via nanoparticle tracking analysis using a NanoSight NS300 (Salisbury, UK). In brief, EV samples were diluted 1:50 in PBS, and 1 mL of sample was injected into the NanoSight at a flow rate of 500 µl/min. Three 60s videos at different sample locations were gathered for size and concentration analysis. Additionally, sEV morphology was also characterized by using negative stain and transmission electron microscopy imaging through the Emory Integrated Electron Microscopy Core.

### Extracellular Vesicle RNA and Protein Isolation

300 µL of serum or 700 µL of EVs were used to isolate RNAs using a Plasma/Serum RNA Purification Mini Kit (Norgen Biotek, Canada) and an Exosomal RNA Isolation Kit (Norgen Biotek, Canada, Cat # 55000) per the manufacturer’s instructions. Quality control of the isolated RNAs was performed using an Agilent Bioanalyzer (TapeStation 4200, Agilent Technologies). miRNA and total RNA library preparation and next-generation sequencing were done by the Emory National Primate Research Center Genomics Core.

Lysates of serum-derived extracellular vesicles (EVs) were prepared utilizing the Easypep MS Sample Prep kit (ThermoFisher, MA, USA). The lysis protocol included sonication in an ice-water bath for a duration of 10 minutes. Following homogenization, protein concentration was quantified using the Rapid Gold BCA Protein Assay Kit (ThermoFisher). The samples were subsequently subjected to reduction, alkylation, and enzymatic digestion to generate peptides for analysis. Chromatographic separation was conducted using a Dionex RSLCnano system. Digested peptides were loaded onto a Waters CSH column (C18 resin, 150 mm length) and eluted at a flow rate of 1.5 µL/min. The elution profile utilized a 55-minute gradient increasing from 1% to 99% Buffer B. The mobile phases were composed of 0.1% formic acid in water (Buffer A) and 0.1% formic acid in 80% acetonitrile (Buffer B). Data acquisition was performed on a Q-Exactive HFX mass spectrometer (ThermoFisher) operating in positive ion mode, utilizing a data-dependent acquisition (DDA) strategy with a Top 20 method. Raw spectral data were processed using MaxQuant software (v2.1.3.0). Peptide identification was performed by searching against a Uniprot sheep (*Ovis aries*) reference database containing 23,110 target sequences. The search parameters defined carbamidomethylation of cysteine (+57.0215 Da) as a fixed modification. Variable modifications included methionine oxidation (+15.9949 Da) and protein N-terminal acetylation (+42.0106 Da), with a maximum of five modifications permitted per peptide.

### Proteomic Data Processing, Differential Enrichment, and Loading Analysis

Raw protein intensity values were first processed to address missing data and systematic variance. Missing values were imputed using K-Nearest Neighbors. Following imputation, the dataset was normalized using Variance Stabilizing Normalization (VSN) via the NormalyzerDE R package^25^. To identify proteins with significant changes in abundance between the single ventricle and control physiologies, we performed differential enrichment analysis using the linear modeling features within NormalyzerDE and limma^26^. Proteins were considered differentially enriched if they met a statistical threshold of an absolute log_2_ fold change ≥ 0.58 and an adjusted significance threshold of -log_10_(P-value) ≥ 1.3.

Unsupervised analysis was conducted to assess global proteomic trends. Principal Component Analysis (PCA) was performed using the prcomp function with centered and scaled data to visualize sample separation based on surgical group and tissue origin. To identify the individual proteins most responsible for the separation between Control and Fontan groups, PC1 loading scores were extracted from the PCA rotation matrix. The top ten proteins with the highest positive loading scores (associated with Fontan enrichment) and the top ten with the most negative loading scores (associated with Control enrichment) were selected and visualized using a diverging bar plot. Hierarchical clustering was generated using the pheatmap package utilizing “Manhattan” distance and the “ward.D” clustering algorithm to visualize expression patterns across samples^27^. To functionally annotate the proteomic changes associated with Fontan physiology, we performed pathway enrichment analysis on differentially enriched proteins using g:Profiler through the gprofiler2 R package^28^. Fontan-enriched proteins, Control-enriched proteins, and all differentially enriched proteins were analyzed separately using *Ovis aries* as the reference organism. Gene Ontology Biological Process, Gene Ontology Molecular Function, Gene Ontology Cellular Component, and Reactome pathway databases were queried. Enrichment results were corrected for multiple hypothesis testing using the false discovery rate method, and terms with adjusted P-values < 0.05 were considered significant. Enrichment scores were calculated as the ratio between the observed query-term overlap and the expected overlap based on the effective domain size. The top enriched terms were visualized using bubble plots in which point size represented the number of overlapping proteins and fill intensity represented -log_10_(P-value).

### Small RNA Sequencing Alignment and Quantification

Raw sequencing reads were processed using the TrimGalore wrapper package to remove adapter sequences and low-quality bases^29^. To ensure high-quality input data, reads with an average quality score below 20 or a read length of less than 18 nucleotides were discarded. The cleaned reads were subsequently aligned to the *Ovis aries* reference genome using the Bowtie short read aligner. Alignment parameters were configured to ensure high specificity, allowing for zero mismatches in the seed region (-n 0) and suppressing reads with more than five valid alignments (-m 5) to reduce multi-mapping ambiguity. The resulting alignments were quantified against the reference annotation to generate count matrices for downstream differential expression analysis.

### Transcriptomic Data Normalization and Differential Expression

Raw count data from RNA sequencing were processed using the edgeR and limma packages^26,30^. Poorly expressed genes were filtered out to reduce mean-variance trend bias. The remaining reads were normalized using the Trimmed Mean of M-values (TMM) method to account for library size differences. To correct batch effects identified during exploratory data analysis, we applied the ComBat function from the sva package on the Log2-counts per million (LogCPM) transformed data^31^.

Differential gene expression analysis was performed using the voom transformation to estimate the mean-variance relationship of the log-counts. To account for the correlation between repeated measurements on the same experimental subjects, we utilized the duplicateCorrelation function twice within the limma linear modeling framework. This approach treated the surgical cohort as a blocking factor. Empirical Bayes smoothing was applied to the standard errors to calculate moderated t-statistics. Transcripts were defined as differentially expressed based on an absolute log_2_ fold change ≥ 1.0 and an adjusted significance threshold of -log_10_(P-value) ≥ 1.3.

Similar to the proteomic dataset, principal component analysis and hierarchical clustering was performed using the prcomp and pheatmap package, respectively, to identify global transcriptomic trends^27^. Hierarchical clustering was generated utilizing “Manhattan” distance and the “ward.D” clustering algorithm to visualize expression patterns across samples. For miRNA pathway analysis, Target genes from differentially expressed miRNAs were identified using miRTarBase^32^. Targets were defined as those having been validated by at least three different experimental methods. Target genes were input on Metascape as a multiple gene list^33^. Only Reactome pathways were considered for downstream analysis.

To evaluate longitudinal behavior of candidate small RNA biomarkers, batch-corrected LogCPM values were merged with sample metadata containing animal identity, operative status, timepoint, and fibrosis classification. Timepoints were converted to numeric months post-surgery, with pre-operative and 0-month samples assigned a value of 0. Longitudinal trajectories were generated for the small RNAs shared between pre-operative and non-fibrotic post-operative samples, as well as for the small RNAs included in the final risk scoring panel. Individual animal trajectories were plotted as repeated-measures line plots, and group-level trends were visualized using mean trajectories stratified by fibrosis classification.

To visualize experimentally supported regulatory relationships between scoring miRNAs and fibrosis-associated pathways, we constructed a miRNA-target-pathway network. miRNA-target interactions were queried from miRTarBase, and targets were retained if they had evidence from at least three independent experimental entries^32^. Because miRTarBase uses arm-specific nomenclature, hsa-miR-31 was mapped to hsa-miR-31-5p for target retrieval. Candidate target genes were intersected with curated gene sets representing Toll-like receptor signaling, NF-κB signaling, Interleukin-17 signaling, and TGF-β/fibrosis signaling. Network visualization was performed using igraph, tidygraph, and ggraph, with nodes representing small RNAs, target genes, and pathway categories.

### Feature Selection via LASSO and Elastic Net

To reduce the dimensionality of the omics datasets and identify robust predictors of liver stiffness, we employed regularized regression models. We utilized the glmnet package within the caret framework to fit Elastic Net and LASSO models^34^. The optimal regularization parameter lambda was determined using Leave-One-Out Cross-Validation (LOOCV) to minimize the Root Mean Squared Error (RMSE). For transcriptomic data, an alpha of 0.1 was selected to balance L1 and L2 regularization. For proteomic data, a LASSO regression (alpha = 1.0) was applied. Features with non-zero coefficients at the optimal lambda value were extracted as variables of importance. Model performance and stability were evaluated by visualizing the coefficient paths and Mean Squared Error across the range of lambda values. To assess redundancy among selected transcriptomic predictors, pairwise Spearman correlation coefficients were calculated across the Elastic Net-selected small RNA features. Correlation structure was visualized using hierarchically clustered heatmaps and coefficient-labeled correlation matrices.

### Development and Evaluation of the Fontan Extracellular Vesicle Score (FES)

We developed a risk stratification scoring system using Ordinal Logistic Regression (OLR) to predict the severity of Fontan-Associated Liver Disease (FALD). The outcome variable was defined as liver stiffness measured by elastography and categorized into three ordinal levels: Stable (≤ 1.6 m/s), Moderate (> 1.6 and ≤ 2.4 m/s), and Severe (> 2.4 m/s). Continuous predictors identified during the feature selection step were discretized into bins to facilitate a point-based scoring system. Cut-off points for binning were determined based on quantile distribution. We fitted the OLR model using the *polr* function from the MASS package^35^. The resulting regression coefficients were rounded to the nearest integer to assign point values to each bin. A total risk score was calculated for each subject by summing the points associated with their specific biomarker levels.

To evaluate the generalizability of the scoring system, a held-out test set of 30 samples was randomly drawn from the full dataset. The training-derived cut points and score lookup table were applied to the test data to compute total risk scores. Model discrimination was assessed using Receiver Operating Characteristic (ROC) analysis with two binary classification tasks: (1) Moderate or Severe fibrosis versus Stable, and (2) Severe fibrosis versus all other categories. ROC objects were generated using the pROC R package^36^. To assess classification accuracy, each test subject was assigned a predicted fibrosis category by mapping their total risk score to the probability lookup table. A subject was classified as Severe if the predicted probability of Severe exceeded 50%, as Moderate if the predicted probability of Moderate exceeded 50%, and as Stable otherwise.

### Validation, Sensitivity Analysis, and Comparison with Established Fibrosis Indices

To quantify uncertainty in model performance, we performed bootstrap internal validation of the test cohort. The test dataset was resampled with replacement 1,000 times, and ROC AUC values were recalculated for each resample for both classification tasks: Moderate or severe fibrosis versus stable status, and severe fibrosis versus all other categories. Bootstrap samples lacking both outcome classes were excluded from the corresponding AUC calculation. Median bootstrap AUC values and percentile-based 95% confidence intervals were calculated from the resulting distributions.

To evaluate whether model performance was dependent on the selected elastography thresholds, we performed a cutoff sensitivity analysis. The Stable-to-Moderate boundary varied from 1.4 to 1.8 m/s, and the Moderate-to-Severe boundary was varied from 2.2 to 2.6 m/s. For each cutoff combination, fibrosis categories were reassigned while retaining the original risk score, and ROC AUC values were recalculated for both Moderate-or-Severe and Severe classification tasks. This analysis assessed the robustness of the scoring system to plausible variation in elastography-based disease definitions.

To benchmark the Fontan EV Score (FES) against established non-invasive fibrosis indices, we calculated FIB-4 and APRI scores for matched timepoints. FIB-4 was computed as (Age × AST) / (Platelets × √ALT), where age was derived from date of birth to the date of the blood draw, AST and ALT were measured in U/L. APRI was calculated as ((AST / AST-ULN) / Platelets) × 100, with the upper limit of normal (ULN) for AST set to 80 U/L based on ovine reference ranges^8,9^. Each test-set sample was matched to the closest available FIB-4/APRI blood draw by first identifying the sample directory entry with the nearest rounded month to the elastography timepoint and then selecting the FIB-4/APRI measurement closest in calendar date to that directory entry. Only matches within a 30-day window were retained. Because FIB-4 and APRI are fixed-formula indices not fitted to the data, their ROC performance was evaluated on the full matched dataset to maximize statistical power, whereas the FES was evaluated exclusively on the held-out test set to prevent inflated performance estimates. ROC AUC values were calculated for both classification tasks and compared across all three indices. Additionally, to contextualize the added value of the multi-analyte FES relative to its individual components, ROC curves were generated for each of the eleven small RNA features independently on the held-out test set. The direction of the ROC predictor was assigned based on the sign of the point value in the scoring system: features with positive points (disease-associated) used an ascending direction, while features with negative points (protective) used a descending direction.

### Liver Organoid Culture and TGF-β-Induced Fibrosis Validation

Induced pluripotent stem cell line CBiPSC6.2 (Gibco, Cat# A18945) was a kind gift from Dr. Gail Besner (Nationwide Children’s Hospital, Columbus OH) and was maintained in Essential 8 medium (Fisher, Cat# A1517001). CBiPSC6.2 was used for differentiation, following a protocol reported previously^37^. Briefly, CBiPSC6.2 at 85-90% confluency was changed with the following media for definitive endoderm (DE) differentiation. Day 1: RPMI supplemented with 50 ng/ml BMP4 and 100 ng/ml Activin A; Day 2: RPMI with100 ng/ml Activin A and 0.2% Knockout serum replacement (KSR, Gibco, Cat# A3181502); Day 3: RPMI containing 100 ng/ml Activin A and 2% KSR; Day 4-6: Advanced DMEM/F12 (Gibco, Cat# 12634) containing 1x B27 (Gibco, Cat# 17504), 1xN2 (Gibco, Cat# 17502), 500ng/mL recombinant human fibroblast growth factor-4 (FGF-4, SinoBiological, Cat# 16043-HNAE), and 3 μM CHIR99021 (MedChemExpress, Cat# HY-10182). On Day 7, DE cells were trypsinized and 750,000 cells were mixed with Matrigel, which was then, aliquoted to form Matrigel domes in a 24-well plate. Advanced DMEM/F12 supplemented with 1x B27,1xN2, 1xPen/Strep, 5 ng/mL FGF2 (SinoBiological, Cat# 10014-HNAE), 10ng/mL VEGF (SinoBiological, Cat#11066-HNAB), 20ng/mL EGF (SinoBiological, Cat# 10605-HNAE), 3 μM ChiR99021, 0.5 μM A83-01 (SinoBiological, Cat# A09-900), and 50 μg/mL ascorbic acid (Thermo scientific, Cat# A15613.36) was used from Day 7 to Day 10. Advanced DMEM/F12 supplemented with 1x B27,1xN2, 1xPen/Strep, and 2 μM retinoic acid (RA, Sigma, Cat# R2625) was then used for Day 11 to Day 14. After Day 15, hepatocyte culture medium (HCM, Lonza, Cat# CC-3198) with 100 nM dexamethasone (Sigma, Cat#D4902), 20ng/mL recombinant human oncostatin M (SinoBiological, Cat# 10452-HNAH), and 10 ng/mL recombinant human hepatocyte growth factor (HGF, SinoBiological, Cat# 10463-HNAS) was used to Day 20 at which point HLOs were collected for lysis in Trizol for RNA isolation and quantitative RT-PCR (RT-qPCR), or fixed in 4% paraformaldehyde (PFA) for immunofluorescent assay.

For the HLO fibrogenic model, day 20 HLOs were cultured in a 24-well ultra-low attachment plate (Alkali Scientific, Cat# CFP24) and stimulated with 20ng/ml TGFβ1 (R&D, Cat# 7754-BH-005/CF) for 3 days. HLOs were then collected for RNA isolation or immunofluorescent assay.

### HLO Immunofluorescent assay

HLOs were fixed with 4% PFA at room temperature for 1 h, followed by permeabilization/blocking in DPBS containing 1% BSA, 0.4% Triton X100, and 10% FBS for 1 h. HLOs were incubated with primary antibodies to HNF4α (Thermo Fisher Scientific, Cat# MA1-199), Albumin (R&D, Cat# MAB1455), Vimentin (CellSignal, Cat# 5741), COL1A1 (CellSignal, Cat# 72026S), ACTA2 (α-SMA, Fisher, Cat# MA1-06110), CD31 (CellSignal, Cat# 3528T), CK7 (CellSignal, Cat# 4465T), or CD11b (BD Biosciences, Cat# MAB1124) overnight at 4°C. The next day, HLOs were incubated with goat anti-mouse or -rabbit secondary antibodies conjugated with Alexa Fluor 488 or 568, followed by 4,6-diamidino-2-phenylindole (DAPI, Thermo Scientific, cat# 62248) staining. Imaging was performed using a confocal microscope (Zeiss LSM 800).

### HLO RNA isolation and RT-qPCR

Total HLO RNAs were extracted using miRNeasy mini kits (Qiagen, Germantown, MD, USA). RNA was firstly reverse transcribed into complementary DNA (cDNA) with iScript cDNA Synthesis (Bio-Rad, Cat# 1708891), and cDNA was then used as template for qPCR with iTaq Universal SYBR Green Supermix (Bio-Rad, Cat# 1725122). Primers are shown in Supplementary Table 1. HLO RNA was used for reverse transcription for miRNA detection via miRCURY LNA RT kit (Qiagen, Cat# 339340), and the cDNA was then used for real-time PCR using proprietary miRCURY LNA miR PCR primers (Cat# 339306, Qiagen). Reactions were run in duplicate and GAPDH or UniSp6 were used as internal controls.

## Results

### Enrollment and Characterization of Serum-Derived Extracellular Vesicles

To investigate the utility of circulating biomarkers in FALD, we enrolled a total of nineteen sheep (n = 19) into the study. This cohort consisted of ten subjects undergoing the Fontan operation (n = 10) and nine surgical sham controls (n = 9). For each subject, we collected blood serum samples and performed liver elastography measurements at both pre-operative (Pre-Op) and post-operative (Post-Op) time points in approximately one-month intervals. In our initial analysis, we also grouped samples into three groups based on the degree of their fibrosis: Pre-Op, Negative (samples that were not fibrotic between 0 – six months), and Fibrotic (samples taken after six months). Representative Picrosirius Red-stained and α-SMA liver sections from the Fontan cohort showed progressive collagen deposition and hepatic stellate cell activation, with minimal fibrosis at the pre-operative timepoint, increased collagen accumulation at 2 months post-surgery, and fibrosis at the endpoint (**Figure S1**).

We subsequently performed a comprehensive characterization of small extracellular vesicles (sEVs) isolated from the serum to validate their identity based on morphology, size, and protein composition. Transmission electron microscopy (TEM) was utilized to visualize the structural integrity of the isolated particles. Imaging confirmed that particles from both Pre-Op and Post-Op samples exhibited the canonical globular morphology characteristic of EVs (**Figure 1a**). We further assessed the size distribution of the isolated particles using nanoparticle tracking analysis (NTA). The analysis revealed a mean particle size of 138 ± 9.7nm for Pre-Op EVs and 98 ± 11.2 nm for Post-Op EVs (**Figure 1b**). Finally, we validated the proteomic profile of the vesicles using mass spectrometry. Both sample groups revealed a robust expression of established EV-specific markers: CD81, CD63, CD9, and HSP90AB1 (**Figure 1c**). Collectively, these data confirm the successful isolation of high-purity sEVs from ovine serum suitable for downstream omics analysis.

**Figure 1.**
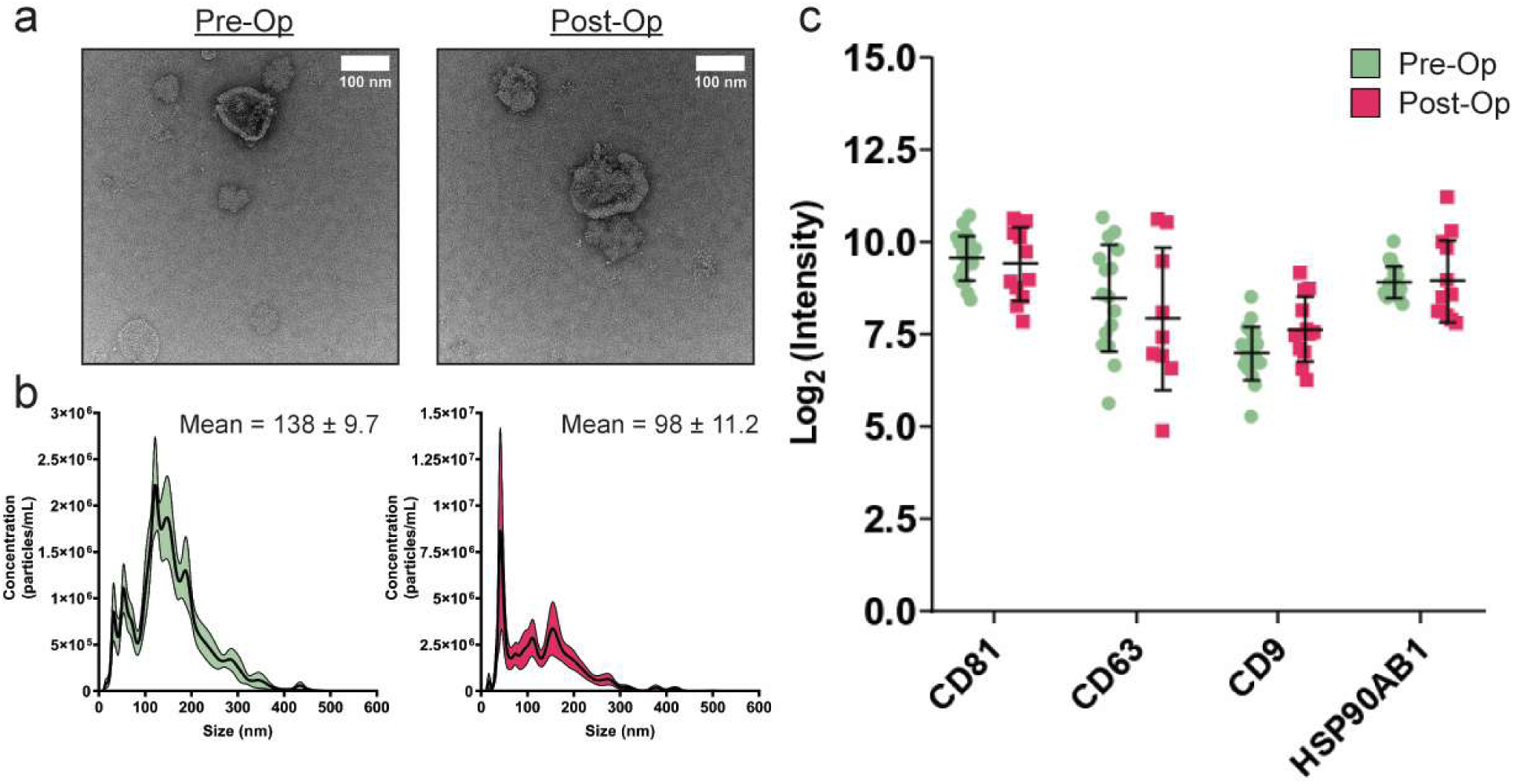
Circulating sEV Characterization. **(a)** Representative negative stain transmission electron microscopy (TEM) images of pre-op and post-op sEVs. The images displayed the canonical cup-shaped morphology characteristic of sEVs and confirmed the successful isolation of intact vesicles. **(b)** Nanoparticle tracking analysis (NTA) of pre-op (mean size = 138 ± 9.7 nm, n = 3) and post-op (mean size = 98 ± 11.2 nm, n = 3) samples. **(c)** Proteomic validation of sEV markers via label-free mass spectrometry. Both pre-op and post-op sEVs showed robust log2 intensity values for canonical sEV markers including the tetraspanins CD81, CD63, and CD9 and the cytosolic marker HSP90AB1.

### Fontan Serum sEVs House Small RNA Cargo Involved in Inflammatory Signaling and a High Hepatic Origin

Following the isolation of serum sEVs, we investigated the transcriptomic content of these vesicles to determine if specific RNA species could serve as biomarkers for hepatic injury. We initially focused on the miRNA content and performed an unsupervised analysis to distinguish between the pre-operative and post-operative groups. We utilized Principal Component Analysis (PCA) and hierarchical clustering to visualize potential group separation. A PCA of the top 50 most variable miRNAs demonstrated no distinct separation between the pre-op and post-op groups (**Figure 2a**). We further compared combinations of Principal Components 1, 2, and 3. All three combinations failed to accurately separate the pre-operative and post-operative groups or distinguish between the pre-operative, negative, and fibrosis cohorts (**Figure S2a**). We cross-validated this finding using a hierarchical clustering approach with the top 50 most variable miRNAs and observed significant overlap between all sample groups consistent with the PCA results (**Figure 2b**).

**Figure 2.**
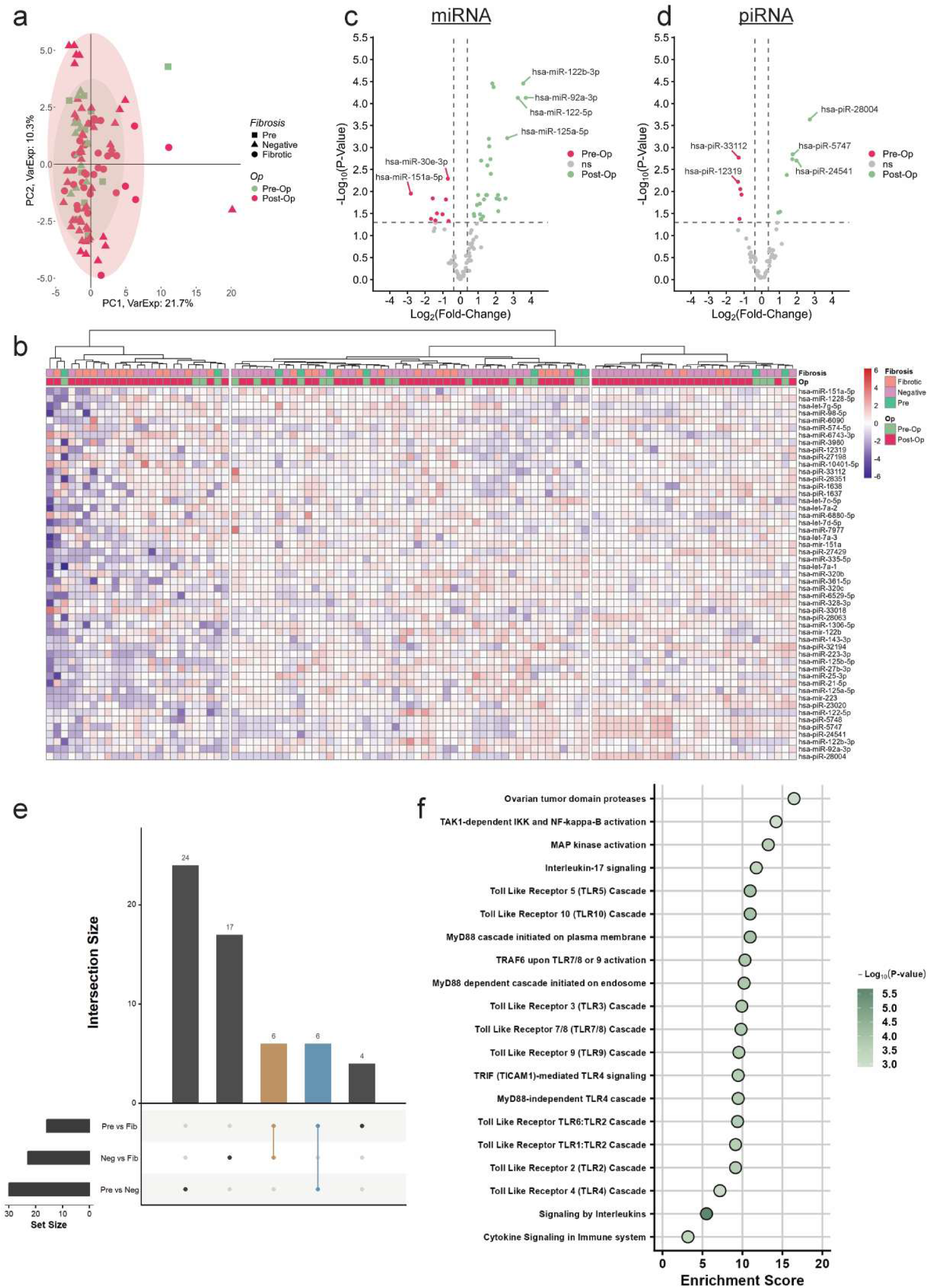
Transcriptomic profiling of circulating serum sEVs in the ovine Fontan model. (**a**) Principal component analysis (PCA) of the top 50 most variable miRNAs showed a lack of distinct global separation between pre-op (green) and post-op (red) groups, as well as between stable and fibrotic cohorts. (**b**) Unsupervised hierarchical clustering of the top 50 miRNAs confirmed significant overlap across all sample groups. (**c**) Volcano plot of miRNA expression highlighting 29 miRNAs that are significantly enriched post-op (green) and 9 miRNAs upregulated pre-op (red). (**d**) Volcano plot of piRNA expression identified 11 piRNAs significantly upregulated in the post-op group and 5 enriched in the pre-op group. (**e**) UpSet plot analysis revealing a core set of six miRNAs that are shared between the pre-operative and negative (non-fibrotic) post-op groups that are subsequently lost or depleted in the fibrotic group. (**f**) Reactome pathway enrichment analysis of target genes for significantly altered miRNAs identified primary involvement in inflammatory signaling axes, including Toll-Like Receptor (TLR) cascades and Interleukin-17 signaling.

We next sought to determine if individual small RNAs could distinguish between the surgical groups despite the lack of group separation. Differential enrichment analysis identified specific miRNA and piRNA species that were significantly altered following the Fontan operation. We identified 29 miRNAs that were significantly enriched in the post-op group and 9 miRNAs that were significantly upregulated in the pre-op group (**Figure 2c & 2d**). Regarding piRNA content, we found 11 piRNAs significantly upregulated in the post-op group while only 5 piRNAs were upregulated in the pre-op group. We subsequently investigated miRNA expression changes associated with the progression of fibrosis by comparing the pre-op, negative, and fibrotic groups. While there was a significant enrichment of nine miRNAs in the comparison between pre-op and fibrotic animals, only one enriched miRNA was identified in the comparison between negative and fibrotic animals (**Figure S2b**). Based on this finding, we investigated the overlap in miRNA expression between the pre-operative and negative groups. An UpSet plot analysis revealed that these groups shared six miRNAs: hsa-miR-320b, hsa-miR-1307, hsa-miR-532-5p, hsa-miR-122b, hsa-miR-6529-5p, and hsa-miR-122b-3p (**Figure 2e**).

To understand the biological implications of these cargo changes, we studied the functional role of the identified miRNAs using pathway analysis. We first generated a list of target genes for the significant miRNAs using the miRTarBase database and performed pathway enrichment analysis via Metascape. This analysis revealed that the miRNA cargo is primarily involved in inflammatory pathways including Toll-Like Receptor and interleukin signaling with the top five enriched pathways being Ovarian tumor domain proteases, TAK1-dependent IKK and NF-kappa-B activation, MAP kinase activation, Interleukin-17 signaling, and the Toll Like Receptor 5 (TLR5) Cascade (**Figure 2f**). Furthermore, a cell type signature analysis of these miRNAs indicated a primary enrichment for Alzarani liver c21 stellate cells (**Figure S2c**).

In summary, while global unsupervised analysis did not reveal broad transcriptomic shifts, differential expression analysis successfully identified a specific subset of inflammatory small RNAs. These miRNAs and piRNAs are associated with hepatic stellate cell activity and inflammatory signaling pathways known to drive liver pathology.

### Proteomic Profiling of Serum sEVs Distinguishes Fontan Circulation from Control Physiology

We next transitioned our focus to profiling the protein content of serum sEVs. For this analysis, we shifted our comparative strategy from a paired pre-operative versus post-operative design to a comparison between Control sheep (n = 9) and Fontan sheep (n = 10) to maximize statistical power given the limited availability of paired samples for proteomic workflows. Concurrently, we sought to elucidate proteomic differences between sEVs derived from serum versus those derived from plasma. In marked contrast to the serum sEV transcriptome, unsupervised analysis of the proteome demonstrated a robust biological signal distinguishing the surgical groups. Hierarchical clustering and Principal Component Analysis (PCA) of the top 50 most variable proteins revealed a clear separation between Control and Fontan samples, with only a single Fontan subject clustering with the Control cohort (**Figure 3a**). The PCA of the sEV proteome displayed a high proportion of variance explained by the first two principal components (**Figure 3b**), and further examination of the component loading vectors indicated that PC1 was the primary driver of the segregation between the Control and Fontan groups (**Figure S3**). We extracted the PC1 loading scores to identify the individual proteins most responsible for this divergence. Proteins driving the Fontan side of PC1 included alpha-2-macroglobulin domain-containing proteins (W5NU00 and W5NUJ7), synaptotagmin-7 (SYT7/W5PUJ6), transferrin receptor protein 1 (TFRC/A0A6P3E4Z4), cathepsin S (CTSS/W5QI70), malic enzyme (ME1/W5PC82), and epithelial/intermediate filament-associated proteins including keratin 6A (KRT6A/W5Q6B8) and keratin 14 (KRT14/W5Q6L8). Conversely, proteins driving the Control side of PC1 included monocyte differentiation antigen CD14 (CD14/W5QJA2), nidogen-1 (NID1/W5P094), fructose-bisphosphate aldolase (ALDOA/A0A6P7DGM3), tenascin (TNC/W5P7G4), triosephosphate isomerase (TPI1/W5P5W9), laminin subunits gamma-1 (LAMC1/W5QD08) and alpha-2 (LAMA2/W5PVY4), phosphoglucomutase-1 (PGM1/W5PJB6), and angiotensinogen (AGT/P20757) (**Figure 3e**).

**Figure 3.**
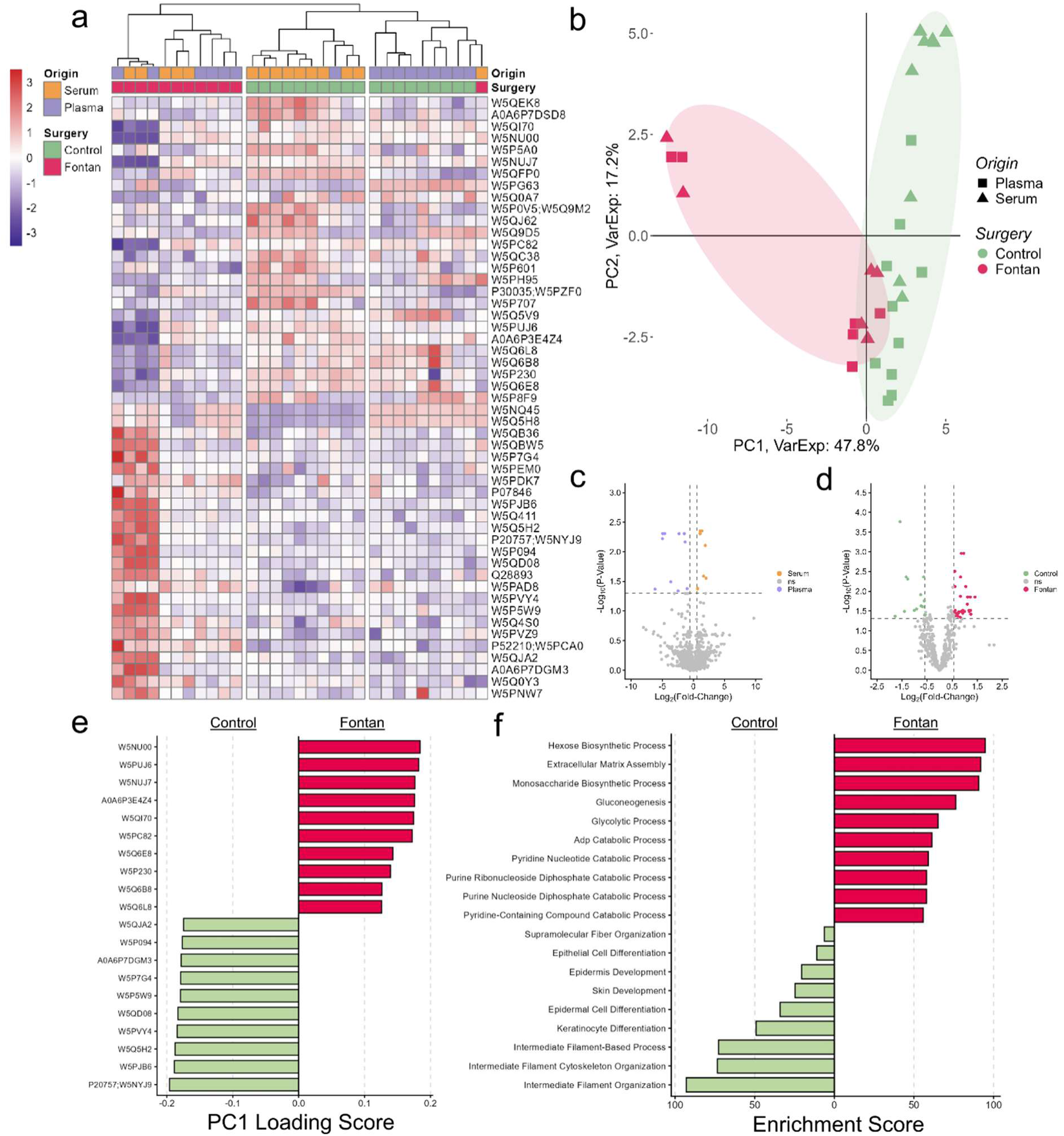
Proteomic profiling of serum sEVs distinguishes Fontan circulation from control physiology. **(a)** Unsupervised hierarchical clustering of the top 50 most variable proteins revealed a clear separation between Control and Fontan samples, with only a single Fontan subject clustering with the Control cohort. **(b)** Principal Component Analysis (PCA) of the sEV proteome showed a high proportion of variance explained by the first two principal components (47.8% and 17.2%, respectively), identifying PC1 as the primary driver of segregation between the groups. **(c)** Differential enrichment analysis comparing serum-derived against plasma-derived sEVs yielded a balanced enrichment profile, with seven proteins significantly enriched in serum sEVs and ten proteins enriched in plasma sEVs. **(d)** Differential enrichment analysis comparing Control and Fontan serum sEVs identified alterations resulting from the Fontan circulation, with 26 proteins significantly enriched in the Fontan group compared to 14 proteins significantly enriched in the Control group. **(e)** PC1 loading score analysis identifying the top 10 Fontan-driving and Control-driving proteins. **(f)** GO Biological Process enrichment with specific pathways named (hexose biosynthesis, ECM assembly, gluconeogenesis for Fontan; intermediate filament organization, keratinocyte differentiation for Control)

We subsequently quantified these differences through differential enrichment analysis. First, we compared serum-derived sEVs against plasma-derived sEVs. This comparison yielded a balanced enrichment profile, with seven proteins significantly enriched in serum sEVs and ten proteins enriched in plasma sEVs (**Figure 3c**). However, the biological comparison between Control and Fontan serum sEVs revealed much more profound alterations. We identified a significantly higher number of differentially regulated proteins in the Fontan circulation, with 26 proteins significantly enriched in the Fontan group compared to 14 proteins significantly enriched in the Control group (**Figure 3d**). To understand the biological implications of this divergence, we performed functional enrichment analysis on these differentially abundant proteins. The Fontan-enriched proteome was highly enriched for metabolic and structural pathways, most notably hexose biosynthetic processes and extracellular matrix assembly, while the Control-enriched proteome was associated with baseline structural maintenance such as intermediate filament organization (**Figure 3f**). Collectively, unlike the small RNA cargo which showed high inter-subject variability, the proteomic cargo of serum sEVs carries a distinct and cohesive molecular signature that strongly differentiates the single-ventricle Fontan physiology from biventricular controls.

### The Fontan EV Score (FES) Stratifies FALD Severity and Outperforms Serologic Indices

Building upon the confirmation that circulating sEV cargo originates from hepatic tissue and distinguishes between surgical physiologies, we sought to leverage these molecular insights to create a non-invasive risk stratification scoring system. The primary objective was to predict the progression of FALD using circulating biomarkers as surrogates for hepatic stiffness. To establish a ground truth for disease severity, we utilized longitudinal liver elastography measurements to bin the sheep into distinct fibrotic categories. We defined these clinical states as Stable (≤ 1.6 m/s), Moderate (> 1.6 & ≤ 2.4 m/s), and Severe (> 2.4 m/s).

We subsequently employed regularized regression techniques to identify the most robust molecular predictors of these fibrotic states. For the transcriptomic profile, Elastic Net regression narrowed the high-dimensional dataset down to 11 key small RNA predictors (miRNAs and piRNAs) (**Figure S4a**). Correlation analysis of these selected features demonstrated low redundancy among the miRNA targets, though a subset of piRNAs clustered together (**Figure 4a**). In parallel, LASSO regression applied to the proteomic dataset identified 12 protein candidates that best predicted fibrosis (**Figure S4b**). However, upon stratifying the available samples into the defined clinical bins, there was a complete absence of matched proteomic data for subjects within the severe fibrosis category. To avoid introducing bias through class imbalance in the high-risk category, we excluded the proteomic features from the final combinatorial model and proceeded with a transcriptomic-only scoring system.

**Figure 4.**
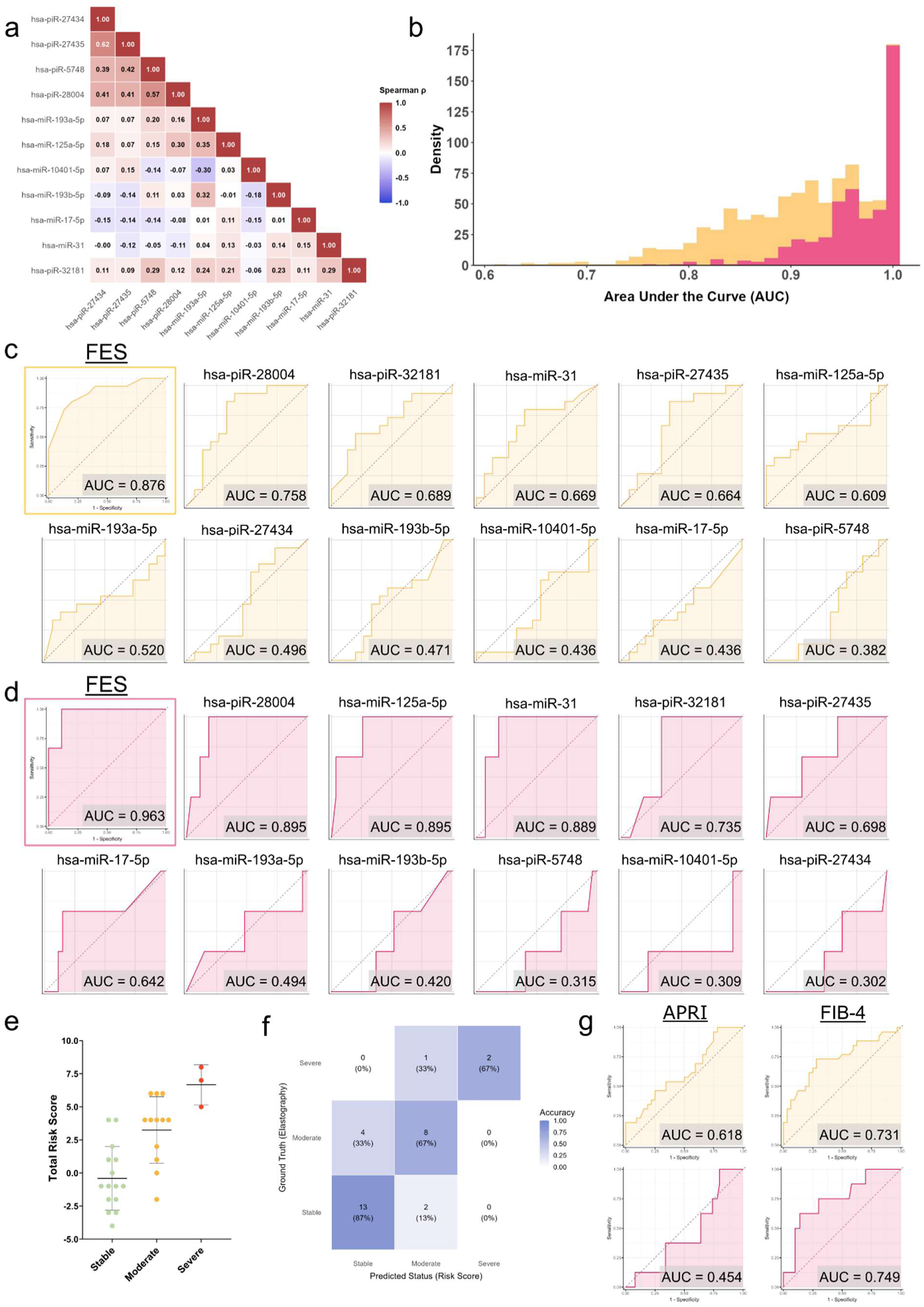
Development of a transcriptomic risk stratification scoring system for Fontan-associated liver disease. **(a)** Spearman correlation matrix of the eleven Elastic Net-selected small RNA features showed low inter-correlation among most predictors, with the strongest association between hsa-piR-27434 and hsa-piR-27435 (ρ = 0.62). **(b)** Bootstrap AUC distributions (1,000 resamples) for Moderate or Severe (yellow) and Severe (pink) classification tasks confirmed stable model performance, with the majority of resampled AUC values exceeding 0.80. **(c)** ROC curves for the combined FES (AUC = 0.876) and each individual small RNA for Moderate or Severe fibrosis detection, with no single feature surpassing the combined score. **(d)** ROC curves for the combined FES (AUC = 0.963) and each individual small RNA for Severe fibrosis detection, where three features individually approached comparable discrimination (AUC > 0.88). **(e)** Total risk score distribution across the held-out test cohort showed distinct separation between Stable, Moderate, and Severe groups. **(f)** Confusion matrix comparing FES-predicted classification against ground truth elastography achieved 87% accuracy for Stable and 67% for both Moderate and Severe categories. **(g)** ROC curves for the established fibrosis indices APRI and FIB-4 for Moderate or Severe (yellow, top) and Severe (pink, bottom) classification, both of which were outperformed by the FES.

We constructed the final risk model using Ordinal Logistic Regression (OLR) to accommodate the graded nature of liver disease progression. The selected predictors were discretized into quantile bins and assigned integer point values based on their regression coefficients (**Table 1**). The model weighted “Time (Months)” heavily (+4 points). Biological predictors associated with increased risk included hsa-piR-5748 (+2 points), hsa-miR-10401-5p, hsa-miR-17-5p, hsa-miR-193b-5p, hsa-miR-31, and hsa-piR-27434 (+1 point each). Conversely, the presence of hsa-miR-125a-5p and hsa-piR-28004 served as protective factors, resulting in a deduction of points (-2 points each) from the total risk score. To translate this additive score into a clinically actionable format, we generated a probability lookup table derived from the OLR model (**Table 2**). This mapping demonstrates a non-linear progression of risk. Subjects with a total score of 0 or lower exhibit a probability of remaining in the Stable category. The transition to intermediate risk occurs rapidly between scores of 3 and 6, where the probability of Moderate disease peaks. Notably, a score of 7 or higher marks a critical threshold where the probability of Severe fibrosis exceeds 60%.

**Table 1.** Risk points for the FALD stratification scoring system. Predictors identified via Elastic Net regularization were assigned integer point values based on their rounded regression coefficients from the ordinal logistic regression model. Continuous abundance of data was discretized using specific scaled count thresholds to facilitate manual scoring. Positive points denoted factors associated with increased fibrosis risk, whereas negative points indicated protective factors.

| Predictor | Threshold<br>(Scaled Count) | Point Value |
| --- | --- | --- |
| Time (Months) | > 3.0 | +4 |
| *hsa-piR-5748* | > 0.193 | +2 |
| *hsa-miR-10401-5p* | > -0.0466 | +1 |
| *hsa-miR-17-5p* | > 0.272 | +1 |
| *hsa-miR-193b-5p* | > 0.0788 | +1 |
| *hsa-miR-31* | > 0.198 | +1 |
| *hsa-piR-27434* | > 0.185 | +1 |
| *hsa-miR-193a-5p* | > 0.215 | -1 |
| *hsa-piR-27435* | > 0.0298 | -1 |
| *hsa-piR-32181* | > 0.357 | -1 |
| *hsa-miR-125a-5p* | > 0.0308 | -2 |
| *hsa-piR-28004* | > 0.0935 | -2 |

**Table 2.** Probability lookup table linking total risk score to disease severity. The cumulative risk score for each subject was correlated with the predicted probability of classification into Stable, Moderate, or Severe fibrosis categories. Thresholds were established to define risk zones, where a score of ≥ 7 indicated a high probability (>60%) of Severe fibrosis, and a score of ≤ 0 predicted a Stable liver with >90% confidence.

| Total Score | P(Stable)(%) | P(Moderate)(%) | P(Severe)(%) | Risk |
| --- | --- | --- | --- | --- |
| -7 | 100 | 0 | 0 | Low |
| -6 | 100 | 0 | 0 | Low |
| -5 | 99.9 | 0.1 | 0 | Low |
| -4 | 99.7 | 0.3 | 0 | Low |
| -3 | 99.3 | 0.7 | 0 | Low |
| -2 | 98.3 | 1.7 | 0.1 | Low |
| -1 | 95.8 | 4 | 0.1 | Low |
| 0 | 90.4 | 9.2 | 0.3 | Low |
| 1 | 79.6 | 19.6 | 0.8 | Low |
| 2 | 61.7 | 36.2 | 2.1 | Low |
| 3 | 40.2 | 54.8 | 5 | Intermediate |
| 4 | 21.9 | 66.7 | 11.4 | Intermediate |
| 5 | 10.4 | 66 | 23.5 | Intermediate |
| 6 | 4.6 | 53.3 | 42.1 | Intermediate |
| 7 | 1.9 | 34.9 | 63.1 | High |
| 8 | 0.8 | 18.8 | 80.4 | High |
| 9 | 0.3 | 8.6 | 91.1 | High |
| 10 | 0.1 | 3.5 | 96.4 | High |
| 11 | 0 | 1.3 | 98.7 | High |

To further characterize the temporal behavior of these biomarkers, we examined the longitudinal trajectories of each scoring panel feature stratified by fibrosis severity. Disease-associated small RNAs generally trended toward elevated expression in moderate and severe subjects relative to the stable baseline, while protective small RNAs trended toward depletion with increasing severity (**Figure S4c & S4d**). However, individual trajectories were highly sporadic, with substantial fluctuations over time within and across subjects.

To evaluate the generalizability of this scoring system, we applied the algorithm to an independent testing dataset (n = 30) and performed bootstrap internal validation. The bootstrap AUC distributions confirmed highly stable model performance across resamples for both classification tasks (**Figure 4b**). When comparing the combined FES to the predictive power of individual small RNAs, the multi-analyte score (AUC = 0.876) outperformed every single feature for the detection of moderate or severe fibrosis (**Figure 4c**). This performance gap was maintained for the detection of severe fibrosis specifically, where the FES achieved an AUC of 0.963, though protective features such as hsa-piR-28004 and hsa-miR-125a-5p showed marked improvement in individual predictive capability for advanced disease (**Figure 4d**).

We further examined the distribution of total risk scores across the testing cohort. The analysis revealed distinct group separation, with score means increasing sequentially from stable to severe, confirming that the scoring system captures the biological gradient of disease progression (**Figure 4e**). Visualized via a confusion matrix, the model demonstrated strong concordance with ground truth elastography, achieving 87% accuracy for stable subjects and correctly classifying 67% of both moderate and severe samples (**Figure 4f**). Finally, we benchmarked the FES against established clinical fibrosis indices, APRI and FIB-4. Evaluated on matched timepoints, the FES substantially outperformed both APRI (AUC = 0.618) and FIB-4 (AUC = 0.731) in detecting moderate or severe fibrosis. This superiority was further magnified when identifying severe fibrosis, where APRI predictive power fell below chance level (AUC = 0.454) and FIB-4 reached an AUC of 0.749, compared to the FES AUC of 0.963 (**Figure 4g**).

In conclusion, this molecular scoring system provides a granular, non-invasive method to stratify FALD risk with significantly greater accuracy than existing serological indices.

### In vitro Validation of FES Biomarkers in a Liver Fibrosis Organoid Model

Having established the predictive performance of the FES, we sought to determine whether the miRNAs in the scoring panel are biologically linked to hepatic fibrogenesis. Pathway analysis of the FES miRNAs revealed enrichment for key fibrosis-associated signaling cascades, including TGF-beta receptor signaling in EMT, TGF-beta receptor signaling activates SMADs, regulation of NF-kappa B signaling, and TAK1-dependent IKK and NF-kappa-B activation (**Figure 5a**). Given this strong association with profibrotic pathways, we sought to functionally validate these miRNAs by testing their response to profibrotic stimulation in a human liver organoid (HLO) model, independent of Fontan hemodynamics. We successfully generated multicellular HLOs from induced pluripotent stem cells (iPSCs) using a 20-day differentiation protocol. Organoids first became apparent on day 11, and by day 20 they exhibited a uniform spherical morphology with diameters of approximately 100 μm (**Figure 5b**). Day 20 HLOs expressed hepatocyte nuclear factor transcripts HNF4A and HNF1B, together with mature hepatocyte markers ALB, AAT, and CK18 (**Figure 5c**). Hepatic stellate cell markers GFAP and PDGFRB were also readily detected (**Figure 5d**). In addition, HLOs expressed the liver sinusoidal endothelial cell marker LYVE1, while the endothelial marker CD31 was detected at levels comparable to those in the definitive endoderm (DE) stage (**Figure 5e**). Cholangiocyte markers CK19 and SOX9 (**Figure 5f**), as well as macrophage markers CD14 and CD68 (**Figure 5g**), were also expressed. Immunofluorescent analysis further confirmed the presence of hepatocyte proteins (HNF4A and ALB), hepatic stellate cell markers (COL1A1 and Vimentin), the macrophage marker CD11b, the endothelial marker CD31, and the cholangiocyte marker CK7 (**Figure 5h**). Together, these findings demonstrate the successful generation of multicellular HLOs containing the major liver cell populations. To establish an *in vitro* liver fibrogenesis model, day 20 HLOs were treated with TGFβ for 3 days (**Figure 5i**). TGFβ treatment markedly increased the protein expression of the fibrogenic markers ACTA2 (α-SMA) and COL1A1 (**Figure 5j**), accompanied by significant upregulation of additional fibrosis-associated transcripts (**Figure 5k**), confirming successful induction of a fibrogenic phenotype. We finally examined the expression of six miRNAs included in the FES biomarker panel. Compared with untreated HLOs, fibrogenic HLOs exhibited marked downregulation of miR-125a-5p, miR-17-5p, and miR-193a-5p, whereas miR-193b-5p was significantly upregulated (**Figure 5l**).

**Figure 5.**
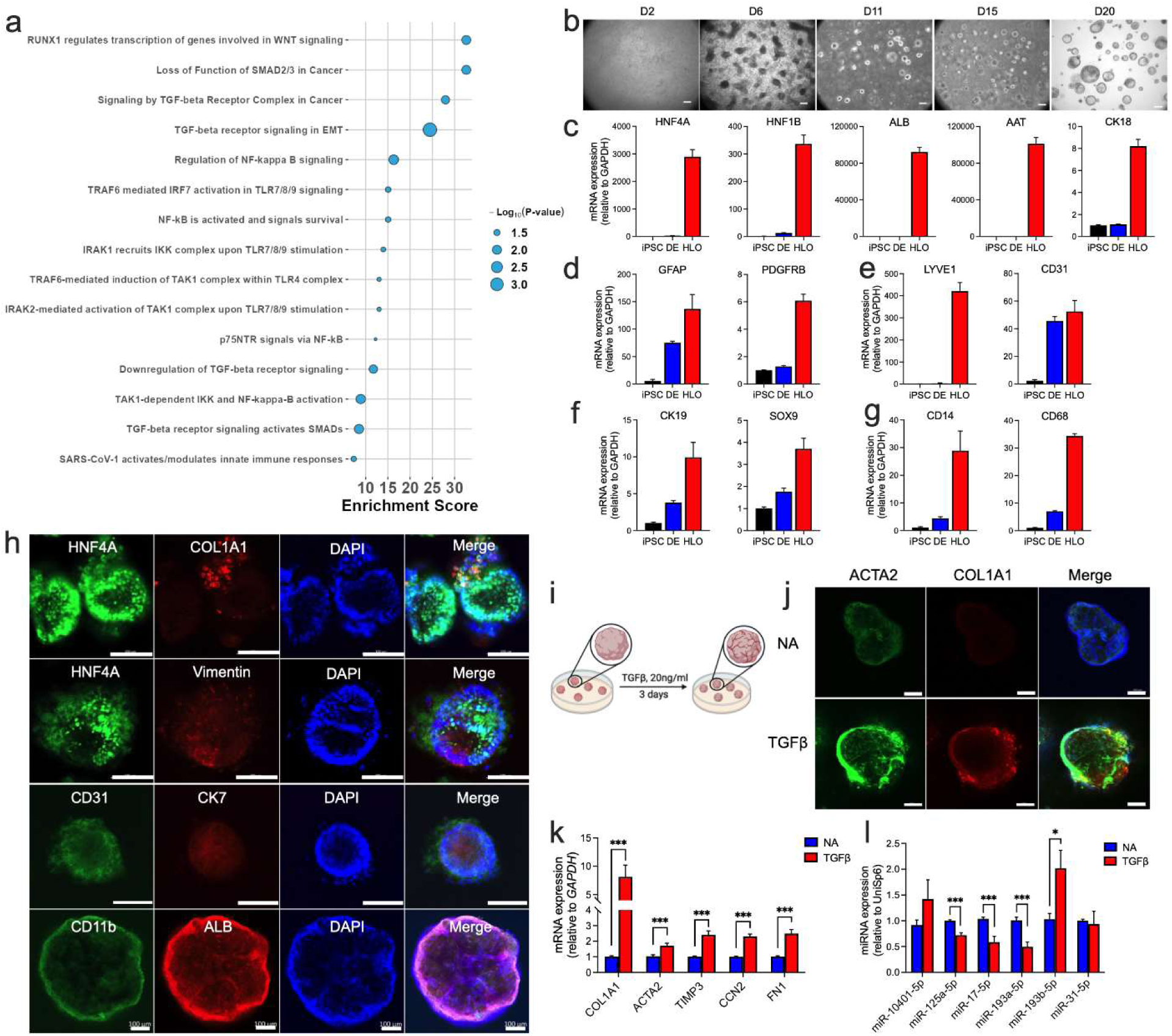
In vitro validation of FES biomarkers in a TGF-β-induced liver organoid model of fibrosis. **(a) (b)** CBiPSC6.2 cells were differentiated into HLOs following a 20-day protocol, cell morphology at different stages were shown. Scale bar=100 μm. HLOs at day 20 were collected, followed by RT-qPCR detection of markers for (**c**) hepatocytes, (**d**) hepatic stellate cells, (**e**) liver sinusoidal endothelial cells, (**f**) cholangiocytes, and (**g**) macrophages. (**h**) Day 20 HLOs were fixed and immunostained for hepatocytes (HNF4A, ALB), hepatic stellate cells (COL1A1, Vimentin), liver sinusoidal endothelial cells (CD31), cholangiocytes (CK7), and macrophages (CD11b). Scale bar=100 μm. (**i**) Model of HLO fibrogenesis: day 20 HLOs were treated with 20 ng/ml TGFβ1 for 3 days. (**j**) Immunostaining of fibrogenic markers (ACTA2 and COL1A1) in HLOs with or without TGFβ1 treatment. (**k**) Transcripts of fibrogenic genes (COL1A1, ACTA2, TIMP3, CCN2, and FN1) were detected by RT-qPCR in HLOs. (**l**) miRNA expressions in HLOs with or without TGFβ1 treatment were detected by RT-qPCR. \**P*<0.05, \*\*\**P*<0.005.

Collectively, these data establish the HLO system as a multicellular human liver model capable of undergoing a TGF-β-induced fibrogenic response. Importantly, the fibrogenic organoids recapitulated directionally consistent changes in several FES-associated miRNAs, including depletion of protective markers such as miR-125a-5p and miR-193a-5p and induction of the risk-associated miR-193b-5p.

## Discussion

Fontan-associated liver disease (FALD) is a nearly universal consequence of the Fontan circulation, driven by chronic venous hypertension and reduced cardiac output^1–3^. Despite its high prevalence, FALD progresses silently and current surveillance tools are poorly suited to detect it. Transaminases are frequently normal because the primary insult is congestive rather than necrotic, GGT elevations are non-specific (Lemmer et al. 2017), and elastography readings are confounded by the elevated central venous pressure inherent to the Fontan physiology^7,38–41^. Consequently, there is a critical need for non-invasive biomarkers that reflect the biological progression of fibrosis at the molecular level.

In this study, we demonstrated that circulating serum extracellular vesicles (sEVs) carry a liver-associated molecular cargo that mirrors pathological changes within the hepatic microenvironment, and we developed the first iteration of a non-invasive scoring system, the Fontan EV Score (FES), for the molecular stratification of FALD severity. Our multi-omic analysis revealed a divergence in how different cargo classes respond to the Fontan circulation. The sEV proteome exhibited robust separation between control and Fontan groups, supported by PCA, PC1 loading analysis, and functional enrichment showing associations with extracellular matrix assembly, metabolic remodeling, and inflammatory reorganization. In contrast, the small RNA transcriptome did not show clear global separation between pre-op and post-op samples in unsupervised analyses, suggesting that while the proteome responds broadly to hemodynamic changes, the transcriptomic landscape may better reflect fibrosis-specific biology.

A key finding was the hepatic origin of the sEV cargo and its involvement in inflammatory signaling. Bioinformatic deconvolution confirmed that a significant proportion of the identified miRNAs originated from hepatic stellate cells, and pathway analysis revealed dysregulation of Toll-Like Receptor cascades, Interleukin-17 signaling, and NF-κB activation in fibrotic subjects. These pathways are well-established drivers of hepatic inflammation and stellate cell activation^42,43^, and their enrichment within circulating sEVs provides evidence that these vesicles report on the inflammatory signaling that drives FALD progression beyond passive congestion.

The addition of the HLO validation experiments strengthens the biological interpretation of the FES transcriptomic panel by testing whether these small RNAs respond directly to fibrotic stimulation in a human liver context. A major limitation of circulating biomarker discovery in the Fontan setting is that molecular signals can reflect multiple overlapping processes, but by using TGF-β-treated HLOs, we were able to evaluate the FES miRNAs in a controlled profibrotic system that is independent of Fontan circulation. The downregulation of miR-125a-5p and miR-193a-5p in fibrogenic HLOs supports their identification as protective features in the FES, while the induction of miR-193b-5p is consistent with its positive contribution to fibrosis risk in the scoring model. The discordant behavior of miR-17-5p, which decreased in the HLO model despite carrying a positive risk weight in the circulating FES, is also informative. This discrepancy may reflect differences between intracellular hepatic expression and circulating EV cargo, or the absence of Fontan-specific hemodynamic and inflammatory cues in the organoid system. Thus, the organoid data should not be interpreted as a complete replication of the *in vivo* sEV signature, but rather as functional support that several FES-associated miRNAs are directly responsive to fibrotic signaling in human liver tissue

One of the most intriguing observations was the overlap in miRNA expression between the pre-op and non-fibrotic post-op groups, with a core set of miRNAs including hsa-miR-320b, hsa-miR-1307, and members of the miR-122b family shared between these stable cohorts but depleted in the fibrotic group. This pattern suggests these miRNAs may function as homeostatic regulators whose loss reflects a breakdown of hepatic homeostasis rather than a response to Fontan hemodynamics^44^. Longitudinal trajectory analysis of the FES features reinforced this concept: risk-associated small RNAs generally trended upward with worsening disease while protective features trended downward, but individual trajectories were highly sporadic. This variability explains why no single small RNA was sufficient as a standalone classifier and why integrating multiple partially informative features into a cumulative score provides a more stable representation of disease state.

Leveraging these insights, we constructed the FES by integrating a single clinical variable, time post-surgery, with circulating small RNA biomarkers. Consistent with clinical consensus, time post-Fontan emerged as the heaviest weighted predictor^3^. The biological functions of the selected miRNAs reinforce the model’s validity: miR-17-5p and miR-31 are established pro-fibrotic mediators linked to Wnt/β-catenin signaling and stellate cell activation^45,46^, while miR-125a-5p is a known protective factor that suppresses TGF-β/Smad signaling^47^. Spearman correlation analysis confirmed limited redundancy among the selected features, supporting the interpretation that the model captures complementary molecular signals. Although our proteomic analysis identified a strong separation between Control and Fontan physiology and LASSO regression selected 12 candidate proteins predictive of fibrosis, proteomic features were ultimately excluded from the current FES. This decision was driven by the absence of matched proteomic data for subjects in the Severe fibrosis category, which would have introduced class imbalance and potential bias into the ordinal logistic regression model. Incorporating proteomic predictors into a future iteration of the FES, once longitudinal proteomic sampling spanning all severity categories is complete, represents a key next step that may further improve discriminative power and biological interpretability.

Beyond the miRNA components, the FES panel also includes five piRNAs, reflecting a recent emergence of piRNAs as circulating biomarkers across disease contexts. Noncoding RNA species, including miRNAs, lncRNAs, and piRNAs, are increasingly recognized as stable and accessible reporters of disease progression when carried by sEVs^48^. piRNAs were originally characterized for their role in transposon silencing in the germline but are now known to be expressed in somatic tissues and have been studied as circulating diagnostic biomarkers in hepatocellular carcinoma and renal cell carcinoma^49^. Of particular relevance to FALD, recent evidence has shown that piR-823 can promote hepatic stellate cell activation through the EIF3B/TGF-β1 axis, directly increasing α-SMA and COL1A1 expression and accelerating liver fibrosis progression^50^. The inclusion of five piRNAs as significant predictors in the Elastic Net model suggests that these transcripts carry independent biological information relevant to FALD that is not captured by miRNAs alone. Further investigation of their mechanistic contributions to hepatic fibrogenesis may reveal novel therapeutic or diagnostic targets specific to the Fontan population.

We validated the FES using ROC analysis, bootstrap resampling, score distribution assessment, and a confusion matrix. The model achieved an AUC of 0.876 for moderate or severe fibrosis and 0.963 for severe fibrosis, with bootstrap resampling confirming stable performance. Classification accuracy was 87% for Stable subjects and 67% for both moderate and severe categories, with misclassifications largely confined to adjacent groups. This performance profile indicates that the FES is effective at ruling out disease in low-risk subjects while retaining utility for identifying patients with deteriorating prognosis.

The comparison against APRI and FIB-4 further illustrates the added value of an EV-based molecular score. Both indices performed modestly in this cohort. However, the FES achieved substantially higher AUCs for both classification tasks, suggesting that circulating sEV cargo provides a more direct molecular readout of hepatic injury. These results position the FES not as a replacement for clinical evaluation but as a complement that adds specificity to existing surveillance strategies.

Our study is not without limitations. The relatively small number of biological replicates (n = 19) may limit extrapolation to the more heterogeneous human Fontan population, and class imbalance in the testing dataset means that the high AUC for severe fibrosis should be interpreted with caution. Our reliance on regularized regression involves trade-offs: Elastic Net and LASSO are effective for dimensionality reduction but may exclude biologically relevant markers from correlated predictor groups. The current FES was developed in an ovine model and will require independent validation in human cohorts before clinical translation. Additionally, although the HLO experiments demonstrate that several FES miRNAs respond to profibrotic stimulation, in vitro organoid models cannot fully recapitulate the combined hemodynamic, inflammatory, and developmental context of the Fontan circulation. It is also important to acknowledge that the current FES relies on elastography as its ground truth for disease severity classification. While we noted in our introduction that elastography readings can be confounded by the elevated central venous pressure inherent to Fontan physiology, it remains the most practical non-invasive measure of hepatic stiffness available for longitudinal monitoring in this model. The present study was retrospective in design, and paired liver biopsies were not available for the cohort used to develop and validate the FES. To address this limitation, a prospective study is currently underway in which ovine Fontan subjects are being followed with both serial elastography measurements and longitudinal liver biopsies. This ongoing effort will enable future iterations of the FES to be calibrated against histologically confirmed fibrosis staging, which will strengthen the biological validity of the severity classifications and help disentangle stiffness attributable to congestion from true collagen deposition. In addition, proteomic features will be incorporated into future versions of the scoring system as longitudinal data spanning all severity categories becomes available.

## Acknowledgements

We gratefully acknowledge the Emory’s EPC Genomics Core for their help with miRNA-sequencing. We also would like to acknowledge the Animal Resources Core at Nationwide Children’s Hospital for the care of the animals used in this study.

## Author Contributions

F. Takaesu conceptualized and designed the study, developed the methodology, performed experiments, conducted the formal analysis, curated the data, and wrote the original draft of the manuscript. X. Li developed the methodology, performed experiments, conducted formal analysis, curated the data, and reviewed and edited the manuscript. J. Kievert developed the methodology, performed experiments, conducted formal analysis, curated the data, and reviewed and edited the manuscript. A. Zhou contributed to formal analysis. S. Kemper, S. Yuhara, S.F. Hussaini, T. Watanabe, J. Matsuda, F. Taha, K. Nelson, J. Zucco, A. Naguib, C. McKee, J. Hill, and S.A. Carrillo contributed to data curation. A. Morrison contributed to data curation and project administration. C.K. Breuer, J.M. Kelly, and D.R. Brigstock provided supervision and project administration. M.E. Davis conceptualized the study, provided supervision, managed project administration, secured funding, and reviewed and edited the manuscript. All authors reviewed and approved the final version of the manuscript.

## Sources of Funding

Funding for this study was supported through the Additional Ventures Cures Collaborative grant, and the American Heart Association and Additional Ventures (AHA/AV) grant (24AVCSASV1276047).

## Disclosures

None.

## Supplemental Material

Figure S1 – S4

Table S1

Major Resources Table

## Non-standard Abbreviations and Acronyms

Pre-Op: Pre-operative
Post-Op: Post-operative
PC: Principal component
AUC: Area under the curve
FES: Fontan extracellular vesicle score

## Notes

### Competing Interest Statement

The authors have declared no competing interest.

